# Dramatic differences in male and female mortality trends for selected European cohorts over ∼20 years

**DOI:** 10.1101/2020.05.23.111971

**Authors:** Jeremy S.C. Clark, Kamila Rydzewska, Konrad Podsiadło, Thierry van de Wetering, Andrzej Ciechanowicz

## Abstract

Longevity is of considerable interest. Collation of recent data after World War II by the Human Mortality Database allowed analyses, previously unattainable, of modal death-ages for sufficient numbers of selected European cohorts. The aim was to track modes and medians/means (≥60 years old (y)) of all-cause mortality for both sexes. The only highest-quality, large-number Lexis data available were analyzed: from nine countries: Denmark, Finland, France, Iceland, Italy, Netherlands, Norway, Sweden and Switzerland; raw-data modes (and medians/means ≥60y, plus thin-plate-spline averages), were analyzed, plus pooled data. Here we show that for cohorts 1880-∼1900 dramatic sex differences existed between death-age changes with all countries except Iceland showing male modal negative trends lasting ∼10-20 years and medians in all countries near-constant or negative lasting ∼10-20 years; whereas females from most countries showed fairly constant positive trends (except Finnish modes and Norwegian medians). For cohorts ∼1900-1919 male and female modal trends were positive (except Dutch and Icelandic cohorts and Finnish females). The net results were that male mortality modes for Danish, Icelandic, Italian, Dutch, Swedish and Norwegian 1919 cohorts were roughly the same as for 1880 cohorts, whereas female death-age modes increased. Results clarify previously knowledge concerning sex differences during this period. Despite improved environment in late adulthood over this period, this did not translate into increased male longevity and earlier events might have sealed their fate, especially in Denmark, Italy, Netherlands, Norway, aand Sweden (and, later, Iceland).

## INTRODUCTION

A lay person’s view of increased longevity often means an expectation that adults have generally been living to increased ages barring early mishap (accidents and/or infectious disease) [1]. The question to be addresssed is whether or how adult longevity has increased for males and females over the recent past for cohorts for which data is now available i.e. for those which have recently become extinct: has this been continuous, intermittent or not at all for either sex ?

To answer this the choice of descriptive parameters is critical. If cohort parameters are available the parameter “life expectancy from birth” is in general not appropriate for this purpose. Ouellette and Bourbeau [2] have stated: “life expectancy at birth for French females is [in 2011] around 85 years….[whereas] the typical age at death has actually been over 90 years in the period life tables since 2006.” Period parameters themselves (from which life expectancies are calculated) cannot give an accurate description of what has happened to death densities at particular times for one particular cohort (a group born at one particular time; in this article during one year; period parameters come from many successive cohorts). The choice to study cohort data, which reflect the true life histories of given birth cohorts, is therefore based on a desire to know what actually happened to particular cohorts; for which we now have full data (i.e. from birth to extinction), with sufficient highest-quality extinct-cohort data only recently available.

Cohort averages are not available after the period studied and in the period before could be severely distorted directly by war.

It was determined to monitor death-density modes with highly-accurate cohort data from the Human Mortality Database (HMD; www.mortality.org), from all countries with Life-Table Lexis data (curated to the highest quality; these were “selected” in the sense that these were the only countries with such data): Denmark, Finland, France, Iceland, Italy, Netherlands, Norway, Sweden and Switzerland, as well as pooled data (or pooled with the two largest countries, Italy and France, removed: “EUM”). There were apparently no other high-quality Lexis cohort life-table data available anywhere for the period studied.

Cohorts were chosen which had sufficient data both sides of the mode to allow spline fits or avoid World-War-II direct-death increases. Modes from raw data, or thin-plate spline interpolation, were analyzed. Additionally, medians and means were calculated for cohort “bulk” deaths, arbitrarily defined as ≥60 years old (y). The three most common measures of location are the mean, median and mode and all can be used to represent “typical” ages of death [3] and the benefits of using these (as opposed to life expectancy) have been extensively discussed by Horiuchi et al. [3] and others.

The major mode is independent of early or late deaths, is therefore a robust measure of location and can be used as a descriptive parameter of adult “bulk” death-ages. For a thorough analysis of both period and cohort modes please see Cheung et al. [4] especially Fig 4 in this reference which shows how cohort major modal ages (referred to in this reference as “late modal ages” and in the present article as “modal ages”) have changed for French, Italian and Swedish cohorts.

Means and medians change according to numbers of earlier (and late) deaths and a limit of ≥60 y was arbitrarily chosen (see Kannisto on use of various median cut-offs [5]). Ages ≥60 y here define the “bulk” of deaths (roughly corresponding to Horiuchi et al.s’ “heap” in [3]).

Instead of using parametric methods (e.g. gompertz, logistic, Weibull, quadratic, normal or skew-t [6]), we followed Ouellette and Bourbeau [7] and Horiuchi et al. [3] in the use of non-parametric splines. Horiuchi et al. [3] discussed the “potential theoretical importance of [the mode] in ageing research” and concluded that use of non-parametric fitting (p-splines) gave “noticeably different” modal trends to those from parametric fits (Gompertz, logistic, Weibull and their Makeham variants). Thin-plate splines can be regarded as more advanced than p-splines in the sense that they produce smooth surfaces infinitely differentiable with an interpretable energy function [8]. They require no manual tuning and visually they seemed to fit well right up to the oldest age. In contrast, with p-splines degrees of freedom are manually adjusted to achieve “best” fits (introducing questions regarding comparisons if different degrees are needed).

Adult mortality is affected by earlier life events, involving factors which affect growth and development of infants, epidemiology, nutrition, salubriousness (sewerage and housing) and schooling [4], as well as access to smoking, red meat and trends in exercise [9]. During the period studied cardiovascular diseases gave the leading cause of world mortality with risk factors from obesity and/or smoking [10]. during this period it was possible that the “bulk” of adult deaths was increasingly determined by lifestyle, including “healthy” (if such a thing exists), mortality. The death distributions of all-cause mortality are therefore likely to become of great interest, especially in high-income countries, as these might start to reflect underlying biological mechanisms of aging.. It was therefore of interest to measure cohort central-tendency location parameters in order to allow future comparisons of timings with life-event parameters.

The study was possible due to recent availability of sufficient cohorts with full data from birth to extinction. The primary question investigated was: how did male and female cohort death-density distributions change over the period of study, using the highest quality data available (from the Human Mortality Database). The main a priori hypothesis was that there were changes in the modal death-ages over time in either sex in the countries analyzed.

The subject of this paper therefore concerned all-cause old-age mortality (including life-style mortality and/or healthy aging) in male and female cohorts born from 1880 to 1919.

## MATERIALS and METHODS

### Data sources

High quality dx data was used from Cohort Life Tables from the Human Mortality Database (HMD; www.mortality.org, RRID:SCR_002370, electronic access from 2015 to 2018, last download 09-11-2018; last checked 12-08-2019), to take advantage of the decisions made by experienced demographers (methods for extracting data, and dx calculations, were double-checked via email communication with HMD). (Further such data is apparently not available.). Data from all available countries: Denmark, Finland, France, Iceland, Italy, Netherlands, Norway, Sweden and Switzerland, were analyzed (source references in Supplemental-file-ORS5).

For each country, all HMD cohort life-table data for one sex (Age interval×Year interval: 1×1) were downloaded into a “country” (Excel 2007; Microsoft, Washington, USA) file; plus Births data. In each country file, HMD columns were removed except: “Year” (which means year of birth in HMD cohort sections; here changed to “Cohort”), “age” and “dx”; and an additional column added: “Year of death” (=“Cohort”+”age”; 110+ as 110). Separate Excel files were created for each cohort (meaning deaths of one sex born in one year), and filtered cohort data from each country file transposed into each cohort file; plus births. In a cohort file, for each country an additional row “Actual (numbers of) deaths” (dx’) was calculated by [dx values×births/100000 (the radix)] (This was checked against the summed “actual deaths” for each country cohort and found to be accurate; for another use of the radix see [4]). dx’ values from each country were summed to give dx’ values for a pooled cohort. Further pooled cohorts with Italy and France removed were created (“EUM”). (All cohort files are combined in “dx_primed_creation.xlsx” Supplemental-files-ORS1,ORS2.) The pooled dx’ data, plus dx’ data for individual countries, were transferred to the columns in “dx_primed_collation” Supplemental-files-ORS3,ORS4.

Data from each cohort is for distinct individuals. Any (rare) missingness procedures were operated by HMD (www.mortality.org).

### Period of study

Cohorts born from 1880-1919 (n=40). To avoid edge effects the early-age and old-age borders were the ages with numbers of deaths 3/4 of numbers dying at the pooled mode (see below, and vertical lines in figure). These borders were independent of the modal age; earlier cohorts were prohibited due to direct excess male deaths around World War II and later cohorts through lack of data. Means and medians were only be calculated for cohorts 1880 to 1904 because of lack of old age data for later cohorts. As well as individual country and pooled analyses from nine countries, the entire study was also carried out with data pooled from seven countries (without Italy and France = “EUM”).

### Interpolation

Interpolation was regarded as an important additional analysis to avoid abrupt changes found in the raw data (with the assumption that people die throughout the year, not just on one particular date !).

### Coding for raw data and spline analyses

A main coding file (Word 2007, Microsoft; Supplemental-file-ORS5_Coding_ INDIVIDUAL_countries) carried R coding to access the data in the Supplemental-files-ORS3,ORS4. Analyses used the R statistical platform version 4.0 (https://cran.r-project.org). A broad plan of the coding follows. For R function stability data zeros were changed to 1 × 10^−2^ and 110+ y data was kept in a separate vector. For one cohort, a thin-plate spline was fitted to the actual numbers of deaths (dx’, Supplemental-files-ORS3,ORS4) from age 40 to the oldest age (ages centered to +0.5y), using the R function *Tps* [fields] with weights 1/dx’ and lambda automatically found by generalized cross-validation. The function *predict* [fields] and 0.01 y grids were used to determine age limits with 3/4 numbers of modal deaths (two vertical lines in figure, see below). Raw modes were identified. Interpolated modal ages were found as follows. A preliminary thin-plate spline mode was found by finding the age with spline maximum (with a tie, the midway point was taken). A fine grid of interpolated age points was created around this age to find the final interpolated thin-plate spline modal age.

Raw medians and means were calculated for each country/sex cohort from age ≥60 y to 110+ (as 110) y. For each cohort the function *integrate* [stats] estimated an interpolated median by integrating successive 0.001 ages until half of the integral for the whole age range was reached, or an interpolated mean by multiplying the integrals by age values and dividing by number of values. Integrals were checked graphically with raw data.

Non-graphical coding in two parts (for females and males) of the coding file is identical with only one part-string difference i.e. “fem” or “MALE”. Each time the main coding file was run it appended 20 columns to the “parameter_results.xlsx” Supplemental-file-ORS6: column modeage2dp = interpolated mode; moderawage = raw mode; rawMedian = raw median; rawMean = raw mean; IntMedian = interpolated median; IntMean = interpolated mean; bulkperctotal = bulk deaths (as % of total deaths); perc76overTotal = % of fraction >76 y; perc60to76 = deaths (%) of fraction ≥60<76 y; perceld0 = deaths (%) of fraction ≥95 y; with prefixes for males (M) or females (F) and country.

Country abbreviations used in this study: Denmark (“De” or “DEN”), Finland (“Fi”, “FIN”), France (“Fr”, “FRA”), Iceland (“Ic”, “ICE”), Italy (“It”, “ITA”), Netherlands (“Ne”, “NET”), Norway (“No”, “NOR”), Sweden (“Swe”, “SWE” or “Sw”) and Switzerland (“Swi”, “SWI” or “Sz”).

### Graphical analyses

Graphs were drawn (R *ggplot* [ggplot2]; Irfanview, www.irfanview.com; Designworks version 3.5, Greenstreet Software, Huntingdon, UK). Loess smoothing (quadratic, span = 0.75) was used, but standard errors are only shown in supplemental figures. For all analyses residuals plots were generated.

### Statistics

Statistics coding, found in the “STATISTICS_CODING and RESULTS” Supplemental-file-ORS7, when run in R read data from Supplemental-file-ORS6, analyzed statistics and produced further graphs. Linear model gradients were used for description only (not for comparison). Initially comparative linear regressions were performed but in many cases assumptions were not met (e.g. heteroscedasticity) and for all male:female comparisons non-parametric Kendall independence tests are shown using Kendall correlation coefficients (analogous to slope comparisons in analysis of covariance, but without assumptions for the latter). These followed conversion from Kendall’s tau to Pearson’s r, as in [11], which is probably conservative. The null hypothesis was that distributions were associated i.e. a significant p value meant that the two distributions were significantly different. Male/female comparison statistical results are printed into Supplemental-file-ORS7, including effect sizes for linear regression and Kendall independence tests (using r = Z/√N). The alpha level for significance was set at 0.05 and all comparisons were two-tailed. Online-Resources are available for this paper and all analyses can be re-run using the coding and data files provided (on Mac, Apple Inc., Cupertino, California, USA or Windows, Microsoft, Redmond, Washington, USA) computers; further details in Supplemental-file-ORS5). The STROBE cohort checklist was used (von Elm E, Altman DG, Egger M, Pocock SJ, Gotzsche PC, Vandenbroucke JP. “The STROBE Statement: guidelines for reporting observational studies”).

## RESULTS

Fig 1a shows that in all countries except Iceland death-age modes had large negative trends at some point between cohorts 1880 and ∼1900 which tended to last from ∼10 to ∼20 years. The timings of these negative downturns varied, with Finnish and French negative trends delayed by roughly 10 years, and the Swiss male trend turning to positive at around cohort 1890. From cohorts ∼1900 to 1919 the Icelandic male modal trend then turned negative (and the Netherlands at a later stage), with other countries showing positive trends. For males in Denmark, Iceland, Italy, Netherlands, Sweden and Norway the overall effect was that the modal death age at cohort 1919 was roughly the same as it was for cohort 1880 (!).

**Fig. 1.**
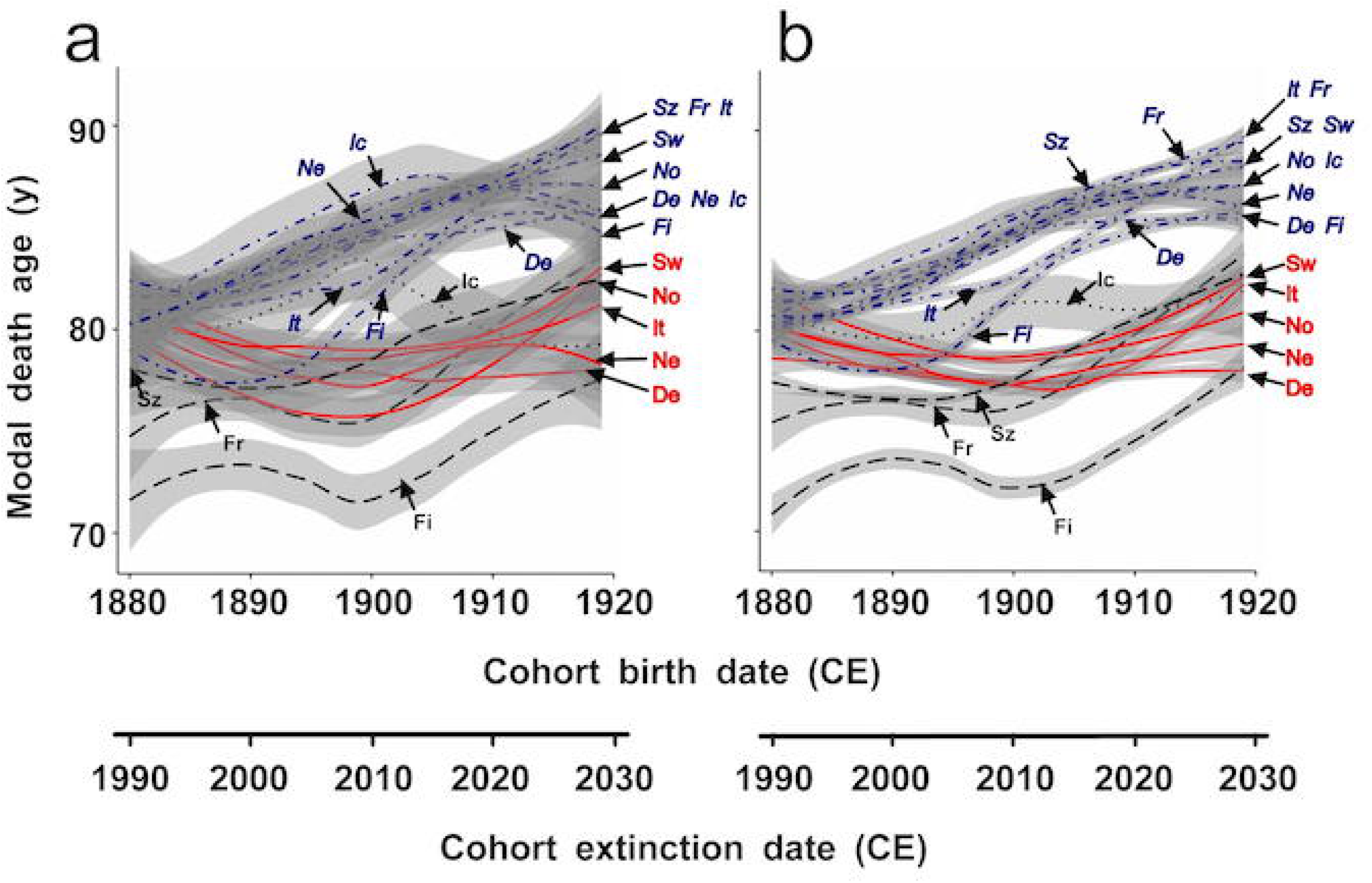
Modal death ages: Individual countries. Female and male cohort modal death ages (years old; y; loess-smoothed; grey areas show standard errors) from (a) raw mortality (dx’) data or (b) interpolated thin-plate splines; from individual countries, versus cohort birth date (common era year; CE) or date cohort defined as extinct (at 110 y). Females (blue dot-dashed lines; italic labels); Males (red or black lines, upright labels): De, Denmark; Fi, Finland; Fr, France; Ic, Iceland; It, Italy; Ne, Netherlands; No, Norway; Sw, Sweden; Sz, Switzerland. Female modes increased, but in five (red solid line) countries, male modes gave negative net modal death age over first 30 years. Source of raw data: Human Mortality Database (2019)

The female trends were dramatically different, with only Finnish females showing negative trend in modal death age between cohorts 1880 and ∼1890, for roughly 10 years, and female modes in all countries showing net positive increases. For cohorts ∼1900-∼1919 female modal trends in some countries (Netherlands, Iceland and Finland) showed negative trends but in Denmark, France, Italy, Norway, Sweden and Switzerland were positive. All female cohorts showed a net overall increase in modal death age between cohorts 1880 and 1919.

The thin-plate spline interpolated modes (Fig 1b) showed similar trends with the sex differences slightly clearer and some timings were slightly different (discussed).

The pooled data (which are naturally weighted by numbers of individuals) provide simple statistical summaries to the above results. Typical pooled mortality density curves are shown in Fig 2, and Fig 2a shows the bump at ∼65 y from direct male World-War-II deaths which deterred use of cohorts before 1880 CE. (Narrow confidence intervals are not shown as these would be hardly visible but can be generated by running Supplemental-file-ORS5.) Pooled modal death ages (i.e. statistical summary graphs) are shown in Fig3 and pooled data without France and Italy are shown in Supplemental-Figs-S1:S3.) All statistical summary graphs simply present the data in a different way but with weighting, and the conclusions drawn are similar to those from individual country graphs (Figs 1 and 4).

**Fig. 2.**
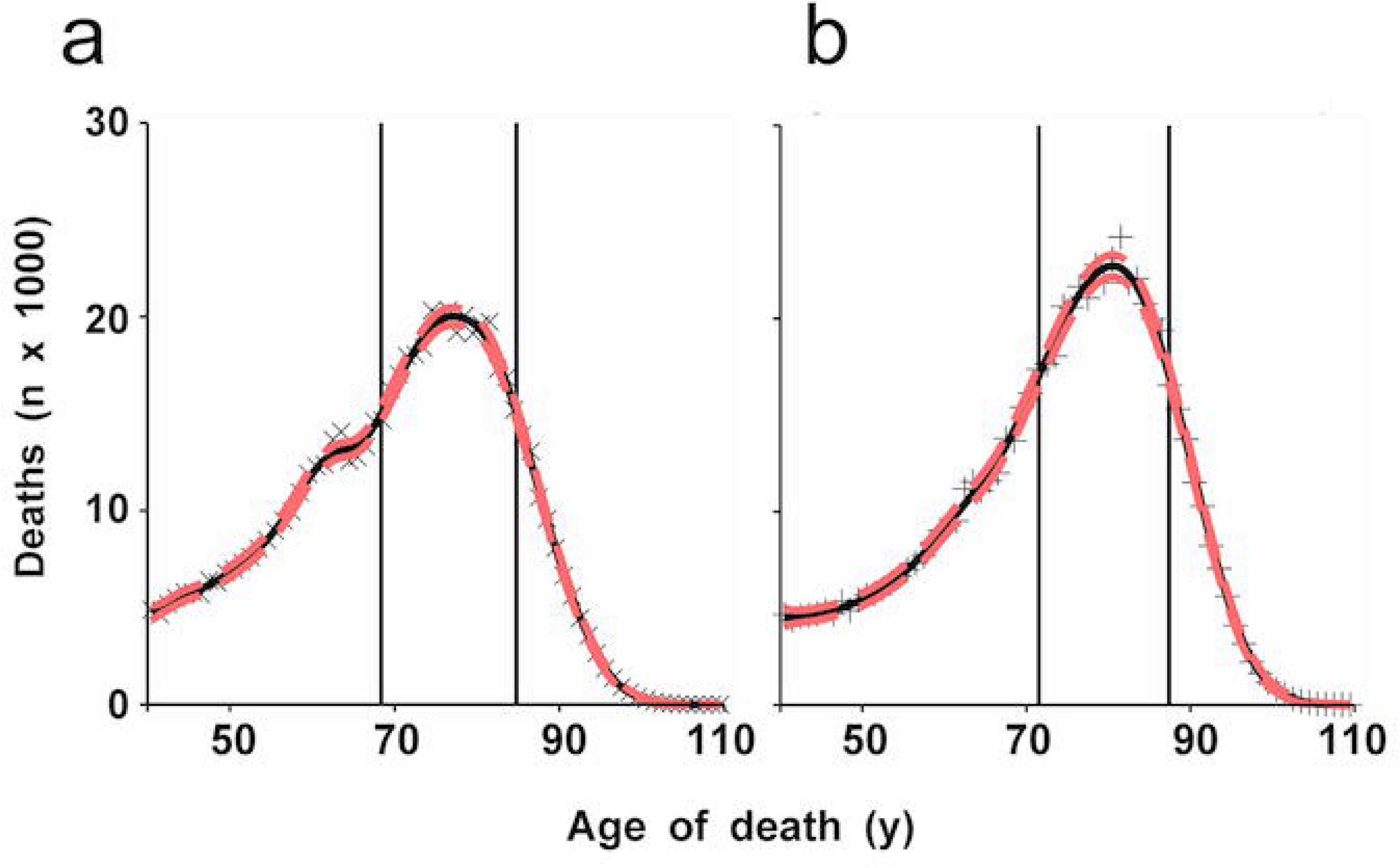
Death (dx’) distribution summary statistics, 1881 CE. Numbers of deaths (estimated from dx) from male (a, x) and female (b, +) pooled cohorts from 1881 common era year (CE) versus age of death (years old; y). Curves show thin-plate spline fits (red dashed lines show narrow 95% confidence intervals which overlay black line fits). Vertical lines show ages with interpolated numbers of deaths 3/4 of numbers dying at the mode. The second-world-war direct-death mortality bump is seen in the male, but not female, cohort. Source of raw data: Human Mortality Database (2019)

Fig 4 shows that in Sweden, Italy, Norway, the Netherlands and Denmark male median or mean death-ages were either near-constant or showed an increase followed by a protracted decrease over the 25 years that medians could be calculated. Icelandic males also showed a decrease after 1890. In France, Switzerland and Finland these male parameters increased (although the medians were near-constant for approximately 10 years during this period, see Fig 4b).

In all countries female bulk medians and means showed increases over 25 years (apart from in Norway during the first 5 years).

Fig 5 shows the overall statistical summaries: male medians constant at 76 y, whereas means increased slightly; female averages increased dramatically over the 25 year period. (Integrated median/mean graphs are similar, see Supplemental-FigS4; and results are also similar if Italian and French data are removed: see Supplemental-Figs-S1:S3) Note the statistical summary graphs simply provide another way of showing the data (but are naturally weighted according to numbers of individuals).

Kendall independence tests showed significant sex differences (p<0.001) for all pooled comparisons (see Figs 3, 5) showing these had changed differently over time. These tests can also be generated for all individual countries by running ORS5_Coding_INDIVIDUAL_countries in R. The Kendall independence tests show whether the male and female changes were associated or not. (Loess-smoothed curves can also be generated, i.e. equivalent to Figs 1 and 4, for all analyses by running Supplemental-file-ORS5 in R and examples are shown in Supplemental-FigS4; the loess standard errors are very small for both males and females, presumably because of the very large datasets.)

**Fig. 3.**
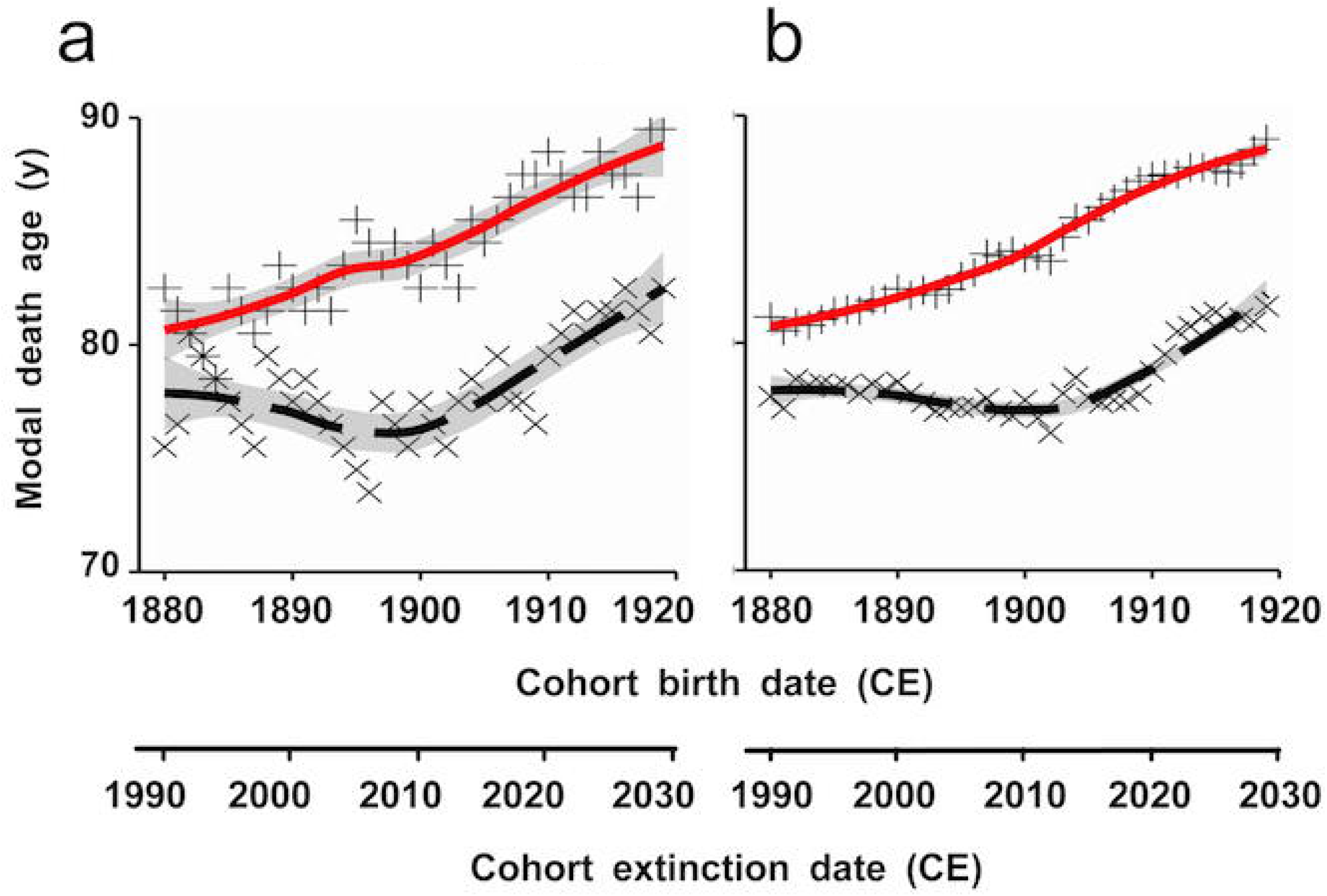
Modal death ages summary statistics. Female (+; solid line: loess, grey areas show standard errors, hardly visible in b) and male (x; dashed line) cohort modal death ages (years old; y) from (a) raw mortality (dx’) data; or (b) interpolated thin-plate splines; versus cohort birth date (common era year; CE) or date cohort defined as extinct (at 110 y). Interpolated pooled modal-death ages show a near-constant increase with female, but not male, cohorts. Source of raw data: Human Mortality Database (2019)

**Fig. 4.**
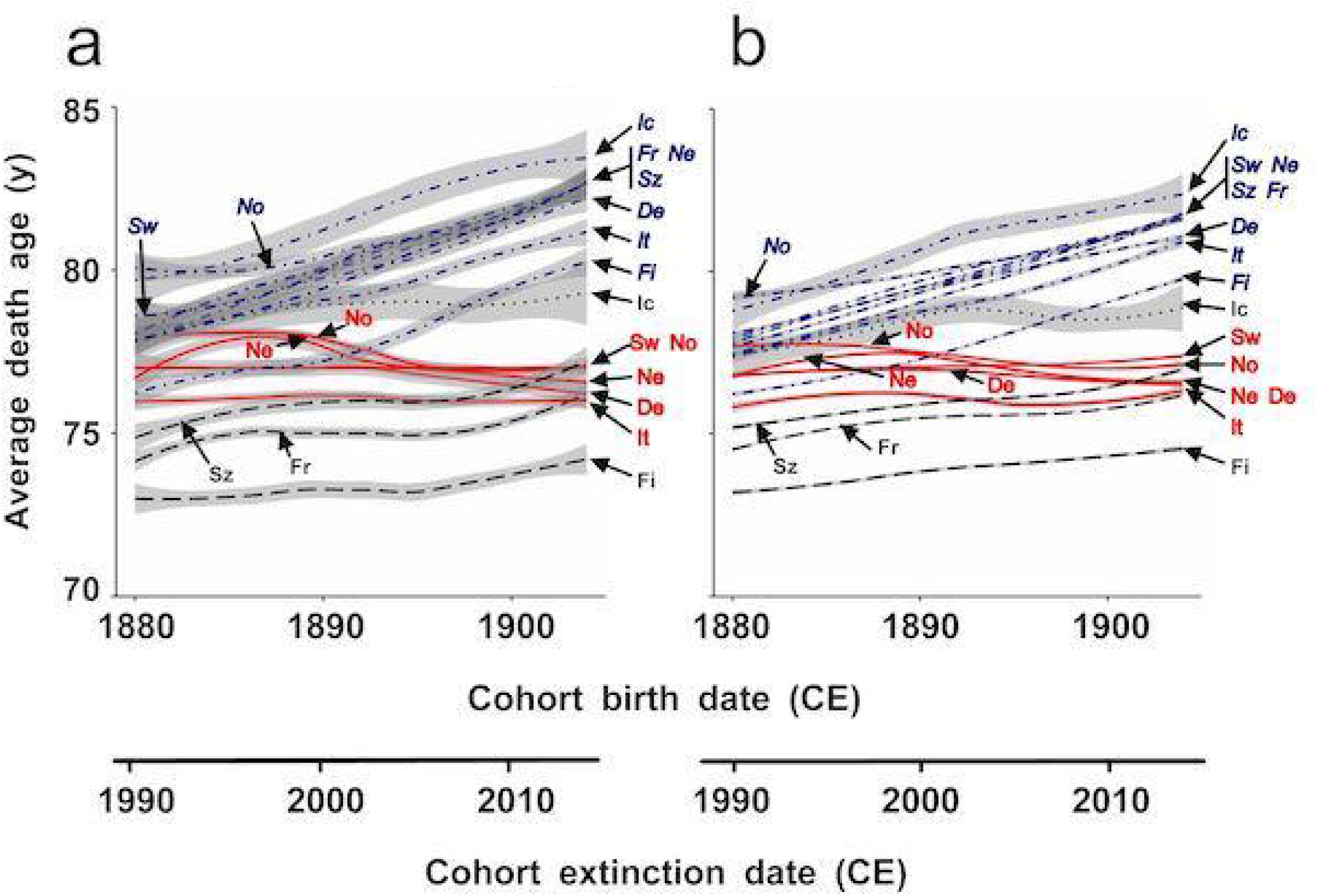
Cohort (a) median, and (b) mean, death ages: Individual countries. Female and male cohort loess-smoothed (grey areas show standard errors) (a) median or (b) mean death ages (years old; y) from individual-country raw mortality (dx’) data, versus cohort birth date (common era year; CE) or date cohort defined as extinct. Females (blue dot-dashed lines; italic labels); Males (red or black lines, upright labels): De, Denmark; Fi, Finland; Fr, France; Ic, Iceland; It, Italy; Ne, Netherlands; No, Norway; Sw, Sweden; Sz, Switzerland. For both averages, female death-ages increased but in five (red solid) countries male death-ages had net decrease or were near-constant over the first 30 years. Source of raw data: Human Mortality Database (2019)

**Fig. 5.**
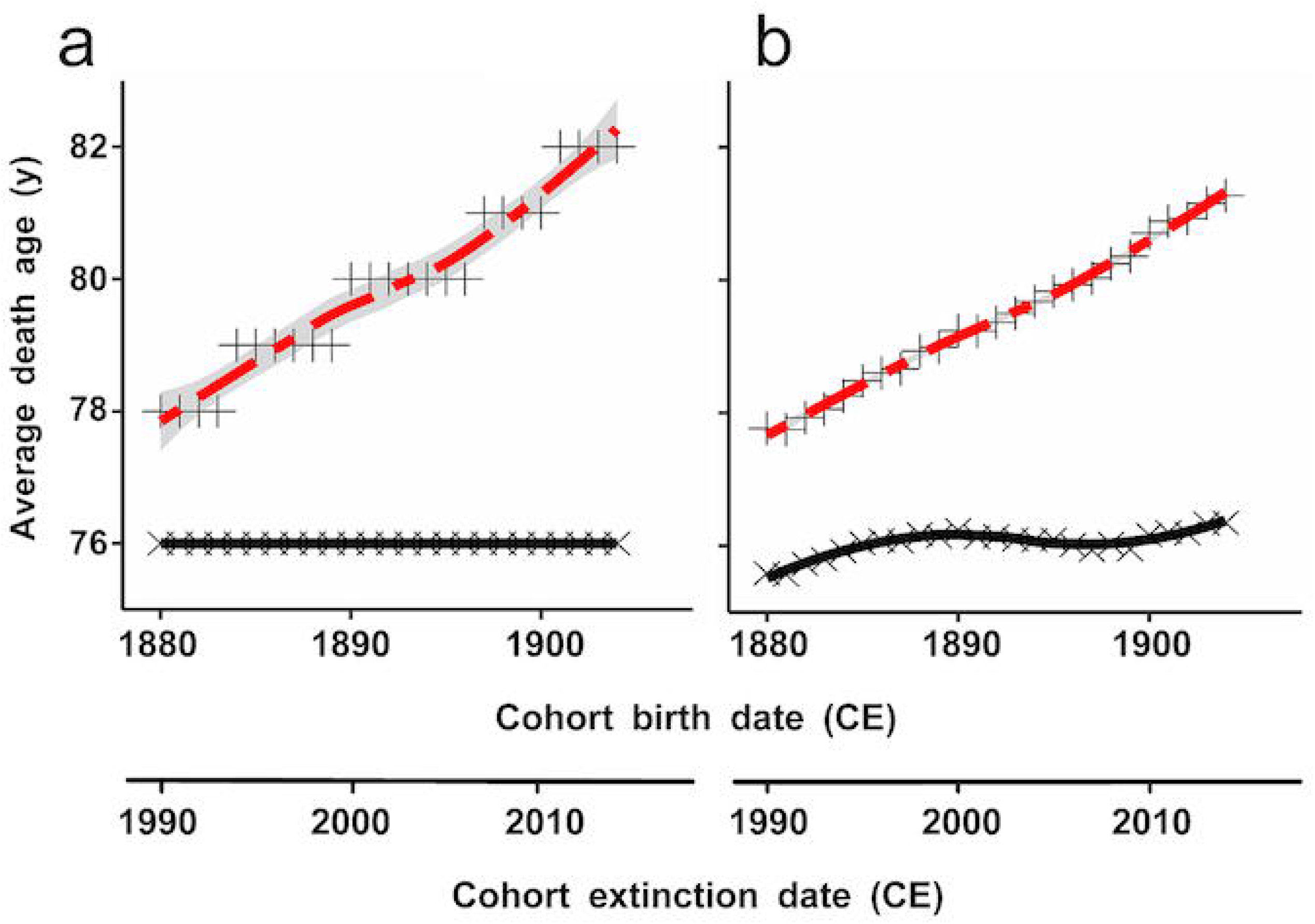
Cohort (a) median, and (b) mean, death ages. Female (+; dashed line) and male (x; solid line) cohort loess-smoothed (grey areas show standard errors, only clearly visible with one line): (a) median or (b) mean death ages (years old; y) from raw mortality (dx’) data, versus cohort birth date (common era year; CE) or date cohort defined as extinct (at 110 y). Raw male median-death ages are exactly constant over this period whereas females’ increase. Source of raw data: Human Mortality Database (2019)

Further statistics including effect sizes and confidence intervals for Kendall independence tests are given in Supplemental-file-ORS7. If Supplemental-file-ORS7 is run in R then results and graphs can be obtained and, in addition, results are also printed into this file. All residuals plots were generated but slight curvatures in the data do not affect the non-parametric statistical comparisons.

## DISCUSSION

### Individual country analyses

The individual country results (Fig 1) are the most important and show dramatic sex differences for all countries in mortality changes over time. male modal averages showed negative trends for at least ∼10 years during the study period and for Denmark, Iceland, Italy, Netherlands, Sweden and Norway the end result was that the modal death age at cohort 1919 was roughly the same as at cohort 1880. In contrast female modal death-ages (Fig 4b) were higher at the end of the study period in all countries. In Fig 3 in Horiuchi et al. [3] similar results can be seen for French males and females (plotted against calendar year, not cohort birth date); and other studies have previously presented modal death-ages [12, 13]. Beltrán-Sánchez et al. [10] analyzed cohorts with birth dates between 1800 to 1935 in 13 countries and stated that “a relatively new demographic phenomenon …. emerged among people born in the late 19th century”, referring to the divergence.

Divergence between male and female modal death-ages in particular countries during the study period has been previously noted by many studies e.g. see [4] (and Fig 7 in this reference; with quadratic fits) in which French and Swedish data are shown for this period (note the x-axis in this Figure shows calendar year, not cohort birth date, and the study concerns mostly period comparisons).

Male medians and means (Fig 4) showed, over the 25 years for which they could be calculated, that Danish, Dutch, Italian, Norwegian and Swedish 1904 cohort values were similar or less than for 1880 cohorts. In these five countries males showed decreasing or near-constant median death-ages over this period (with a hump with Netherland medians resulting in an overall decrease). Net median and mean changes for Finland, France, Iceland and Switzerland were positive, although note that the medians for these countries were near-constant for a considerable period (∼10 years). It is clear from Fig 4a that the male median death-ages in all countries did not change in accordance with female medians, which all showed rapid increases (with Norway female median increase delayed by around 5 years). (Integrated results, shown in Supplemental-FigS4, were similar to the raw results shown in Fig 4.)

Means are highly sensitive to changes in the death-ages of the oldest old and should therefore be treated with caution. Technically conclusions regarding modes (with 40 years of data) are independent of other distributional characteristics whereas the medians (25 years of data) depend on the lower cut-off boundary chosen: here ≥60 y, but are also more robust than means. Conclusions are therefore mostly drawn from the modes which cover 40 years of the study and, to a lesser extent, the medians, which cover the first 25 years.

### Thin-plate splines

Thin-plate splines are not necessary to show the main effects which can be seen quite clearly using raw data. However, thin-plate spline modal results not only showed reduced standard errors around loess lines (run Supplemental files to see this), but also the timings of some modal changes were noticeably different from raw data fits: compare Fig 1a and 1b; for example, the peak for male Icelandic modes is at ∼1898 for raw data, but ∼1902 for thin-plate splines, and the male lines for Netherlands and Denmark differ. It is likely that p-splines would also give different timings but not necessarily exactly as for thin-plate splines, and the latter could be regarded as optimal. The accuracy of the thin-plate spline fits can be seen visually by looking closely at fits to the oldest old up to 110 y in the cohort graphs in Fig 2 and in those seen if the main coding is run from Online-Resource-ORS5: the fits appear to be good in every case, despite the low amounts of data at the oldest ages.

It might be argued that the smoothing of the modes provided by the thin-plate splines is too great but this is not the case: in Fig 3a the male raw modal death ages give iterated downward slopes corresponding to peaks resulting from years with excessive mortality which affected several cohorts at once at different ages. Correct smoothing can also be seen by running the main coding (Supplemental-file-ORS5): the graphs of which will run like a motion picture film, in which such peaks will be seen to move from right to left while the thin-plate spline mode is more stable.

### Study limitations

(1) As Crimmins et al. [14] have indicated, conclusions might well only apply to the countries studied. For many large countries e.g. the USA or China, life-table Lexis data were not available for the period studied, and caution is necessary to draw conclusions from other data types e.g. period data. We could not add data similar to that analyzed because, as far as we know, none exists. (2) Pooled conclusions do not necessarily apply to Europe as a whole, and are only given as summary statistics (The nine countries contributed ∼30% of the non-Russian/non-Ottoman European population [15]). (3) Although conclusions regarding modes are independent of population fraction, medians and means are affected by the fraction analyzed (here ≥60 y) and medians and means would change if another fraction were considered.

### Health, stress and mortality

It is not known why some male average death-ages showed decreases (modes) or were usually near-constant (other averages) in contrast to female parameters, over parts of the study period. Crimmins et al. [14] stated that “our strongest conclusion is that male/female differences in health are highly dependent on historical time and geographic location.” This is certainly true for the countries analyzed and we sincerely hope that downward trends in male mortality parameters are quirks due to historical events which will not be repeated, but this cannot be guaranteed.

It is possible that downward trends in male mortality were predominantly affected by serious male-biased factors which off-set positive late-age factors, for two reasons: (1) during the period in which these males were actually dying (at least after World War II), there were fairly constant increases in gross domestic product per person, health care, diet and general living standards in all countries studied, suggesting that male mortality was either not affected by these improvements or there were factors which seriously offset these; (2) female average death-ages increased and presumably females were exposed to many of the same factors.

Current thinking holds that as mortality due to infectious diseases decreased, then cardiovascular disease, cancers and other chronic diseases became more important factors influencing mortality [14]. Mortality, however, has multivarious causes, and factors affecting infancy, adolescence and young adulthood might well have more influence than is at first apparent. According to Cohen and Oppenheim [16] mortality depends on environmental, behavioural, sanitary, nutritional, and medical conditions as well as public health, social, political and economic organisation, food supplies, education and human values.

Speculation to explain the divergence in male/female mortality rates from 1880 onwards might include:

A. a greater vulnerability of males to cardiovascular diseases and a higher frequency of lethal conditions such as heart disease, stroke, and diabetes [14]. Perhaps most importantly for 50-70 year olds from the cohorts studied, dramatic sex differences were found in UK cardiovascular disease deaths which began ∼1925, with male numbers increasing to ∼1975, but female numbers decreased (see Fig 1 in [17]), and similar trends have been found in other European countries [18]. Quite possibly this increase in male-cardiovascular and other life-style disease frequencies contributed to male longevity stagnation and after this period there was a measured fall in e.g. cardiovascular diseases, which might explain why the male modal death ages in most countries subsequently increased. The genetic basis as to why men might have been more susceptible to chronic diseases such as cardiovascular diseases is well known, with strong sexual dimorphism in aging and survival disadvantage among men likely resulting from a male-specific mitochondrial mutation load, which might also affect (B) and (C) below [14].
B. different behavioural responses to environment, including male increased uptake of smoking [14] (relative to women), and with increased wealth an increase in access to red meat, alcohol and less overall exercise, which all increased the risk of cardiovascular diseases. It is thought that men were “more likely to engage in risky and dangerous behavior and women more likely to engage in health-seeking behavior” [14, 19]. Animal product consumption has been suggested to have significantly contributed to cardiovascular and/or cancerous diseases, and there are hints that they may have had greater negative effects in men than women [20]. Possibly males did not (or were unable to) take advantage of beneficial environmental changes. We can speculate that as prosperity increased, women thrived with better life-style choices, whereas males might have relatively consumed more red meat and alcohol, smoked, and become less active: all risk factors for an earlier death [21].
C. greater susceptibility of males to the long term effects of the 1918 influenza pandemic (colloquially known as the “Spanish Flu”). This hypothesis cannot be discounted as a major determinant yet because of the huge numbers of people infected (it caused acute illness in 25–30 percent of the world’s population [22]) plus the fact that the sex differences in infection rates were large with an age-standardized death-rate difference of 174 per 100 000 [23]. According to Azambuja [24] men of European descent born from 1880 to 1900 (20–40 years old) were preferentially killed because of an unusual immune response and survivors of infection might also have exhibited the same response giving a “primed” later susceptibility to coronary heart disease mortality [24]. While (C) is an interesting theory, and one which could be further investigated by comparing the timing of age-distributional infection rates in individual countries with the graphs in Figs 1b and Supplemental-FigS4, it should be noted that recently some evidence has been gathered against this idea [25]. In this reference Wilson et al. compared >1000 military males who had contracted 1918 influenza with controls and found no differences in long-term survival [25], but as the authors mention this is probably not applicable to whole populations including non-military-personnel. In this regard, as the world is currently in the midst of a Covid-19 pandemic with the SARS-CoV-2 known to give cardiovascular complications [26], lessons learned from this period might be directly relevant to those living today.
D. Although explanations concerning cardiovascular diseases seem persuasive, in any case for the generation studied many had fought in two wars and the deferred effects of war (psychological/injury) might have exacerbated behavioral or other differences. It may be that the cohorts studied show peculiar features with atypical mortality patterns which might reflect, for instance, harmful effects from the two world wars. Such effects can be seen among cohorts which directly participated in a war, or in people who during adolescence suffered from undernourishment due to a war. However, it must be remembered that war-related hypotheses need to account for differences (or lack of them) between countries at war and neutral countries: in particular note that Sweden and Switzerland were neutral throughout both world wars but have different male trends (see Fig 4), and France and Italy were at war in both world wars but also have different trends (see Figs 1 and 4). Denmark, The Netherlands and Norway were neutral throughout World War I, and have overall decreases in average death-ages during this period, which might count against a war-related hypothesis (unless war had positive effects on later mortality).
E. Excess male migration is also a factor which might be considered (with predominantly healthy males migrating). Any theory which purports to explain the presented results will need to explain why the scourge which affected males did so at different times in different countries. It is therefore interesting to note that, as Crimmins et al. [14] have indicated, hypertension levels (a risk factor for cardiovascular diseases) are usually greater for males than females in most countries, but there are several countries where the prevalence of hypertension is higher for women, whereas usually the reverse is the case. At some time after the study period sex differences in mortality changes likely became much less apparent. According to Crimmins et al. [14] “sex differences in disease prevalence and mortality rates may recede” as risks for cardiovascular disease mortality reduce and as men and women behave more similarly. For example, data from the National Health and Nutrition Examination Survey (NHANES) showed that by 2010 there were no sex differences in mean age-specific cardiovascular risk markers at ages >50 years [27], and in the USA and Europe there has been increasing similarity in smoking habits between men and women [28]. The use of optimal parameters will be critical in the analysis of the timing of past events. The timing in modal death-age changes sometimes depended on whether thin-plate splines were used or not; these theoretically providing a more accurate representation. In the future, as cohort parameter data catch up with data collected on health dimensions (which have only been collected since the 1980s with national-level surveys on multiple dimensions of morbidity for large samples of both sexes [1, 14]) it will be the cohort parameters with optimal smoothing functions which will provide the correct insights. From the presented results it might be possible to resolve factors by investigation of relative timings, e.g. of the 1918 influenza pandemic, in different countries, especially for example in comparison with Iceland. We hope that the way in which the data has been presented (which we think is optimal) will encourage further research into these mortality changes - especially relative to the timings of the many factors possibly associated with mortality.

### Notes on pooled analyses

The main results and conclusions do NOT depend on the pooled results, which are simply statistical summaries with natural weighting according to the numbers of individuals. It is easy to see from Figs 1 and 4 that male averages proceeded roughly from left to right and the female averages proceeded roughly upwards showing increases in average death ages. However, it is nice to have statistical summary graphs (Figs 2, 3, 5) as these show the trends rather clearly. Note the following:

1. It might seem strange to pool the results from large and disparate countries such as Italy and France, but it must be remembered that the large and disparate cities of Milan and Naples have already been pooled in the Italian data, and likewise for Paris and Marseille for France. At present we have no idea whether the overall timings in male negative (or near-constant) trends differ or are similar between these four cities. Italy and France contributed approximately one third each to the numbers. If these countries are removed from pooled analyses (see Supplemental-Figs-S1:S3) the shapes of the curves are roughly similar, showing that the distribution shapes are robust.
2. All male data show negative modal trends over some periods (due to unknown causes), and all male median data show periods of near-constancy, and therefore have some similarity.
3. There might be some reluctance to analyse individual country data further, on the grounds that it is not known whether differences in the average parameters are bounded by country borders, or whether, for example, they vary from city to city, plus the fact that some countries, e.g. Iceland, have considerably less data and are therefore more influenced by random fluctuations in regulating factors. The pooled data from nine countries represent the sum total of highest-quality data knowledge concerning the mortality density of these cohort bulks and the large datasets have smooth density distributions (even without splines, see Fig 2; note also that they appear visually to have more regular differential changes), which is less apparent with individual country data.
4. Even if you don’t accept points (1) to (3) it still remains that these are valid statistical summaries.

### Other types of study

Life expectancy at birth could also be compared, but the latter is inappropriate to answer the question raised if cohort parameters are available, and this has not been done (see Introduction; similar arguments apply to life expectancy from age 60 y). As Cheung et al. [4] have stated: “the advantages of using the late modal age at death ……[have been] regularly underlined in the past [15, 29, 30]”.

The present study has used some of the best techniques and only the highest quality data available to calculate cohort death-density modes using raw data or thin-plate splines. Other parameters could be used e.g. mortality rates, but we think dx’ gives clarity.

## Conclusions

In summary, for cohorts 1880-∼1900 dramatic sex differences existed between death-age changes with all countries except Iceland showing male modal negative trends lasting ∼10-20 years and medians in all countries near-constant or with negative trends lasting ∼10-20 years; whereas females from all countries showed fairly constant positive trends (except Finnish modes and Norwegian medians). For cohorts ∼1900-1919 male and female modal trends were positive (except Dutch and Icelandic cohorts and Finnish females). The net results were that male mortality modes for Danish, Icelandic, Italian, Dutch, Swedish and Norwegian 1919 cohorts were roughly the same as for 1880 cohorts, whereas female death-age modes had net increases. There are many possible causes for the sex differences in the timings of mortality changes for cohorts born during this period. Suspicion lies with changes connected with cardiovascular and other “life-style” diseases.

## Supporting information

Supplemental File ORS1

Supplemental File ORS2

Supplemental File ORS3

Supplemental File ORS4

Supplemental File ORS5

Supplemental File ORS6

Supplemental File ORS7

## Acknowledgments

We would like to thank the Human Mortality Database staff for their help and availability of data on their website at www.mortality.org.

## Research Ethics Approval

Not applicable - ethics considerations were previously dealt with by the Human Mortality Database. Data is freely available online

## Consent to participate

Not applicable

## Consent for publication

Not applicable

## Availability of data and material

All data, code and all calculations can be found in the Online Resources and the Human Mortality Database (www.mortality.org)

## Code availability

All data and all calculations can be found in the Online Resources. All residual plots can be generated using Online Resource coding. Correspondence and requests for materials should be addressed to JSCC.

## Authors’ contributions

JSCC contributed to conceptualization/data curation, formal analysis/interpretation, statistical analysis, writing of article; KR, KP, TW to formal analysis/interpretation, statistical analysis, critical revision of article, AC to supervision.

Data source references (required by the Human Mortality Database rules) are found at the end of Supplemental-file-ORS5.

## Supporting Information

**Supplemental files: ORS1_Female_dx_primed_creation**.**xlsx**

Pooling calculations for Pooled Female data from nine countries.

**ORS2_Male_dx_primed_ creation**.**xlsx**

Pooling calculations for Pooled Male data from nine countries.

**ORS3_Female_dx_primed_collation**.**xlsx**

Collated dx’ female data from Pooled data, EUM (= with Italy and France removed) and individual countries.

**ORS4_MALE_dx_primed_collation**.**xlsx**

Collated dx’ male data from Pooled data, EUM (= with Italy and France removed) and individual countries.

**ORS5_Coding_INDIVIDUAL_countries**.**docx**

Main coding file. If all header rows are removed (apart from one) from ORS3 and ORS4 and these saved as .csv, when ORS5 is run in R it will read ORS3 and ORS4 and produce results for chosen country, or for Pooled data or EUM, which will be written to ORS6.

**ORS6_parameter_results**.**xlsx**

Results from running ORS5.

**ORS7_STATISTICS_CODING and RESULTS**.**docx**

Statistics coding file. If all header rows are removed (apart from one) from ORS6 and this saved as .csv, when ORS7 is run in R it will read ORS6 and generate statistics and graphs. Results have been printed into ORS7.

## Supplemental Figure legends

**Supplemental-FigS1 Death (dx’) distribution summary statistics for seven countries, 1881**. Data pooled from all countries except France and Italy (“EUM”). Numbers of deaths (estimated from dx) from male (a, x) and female (b, +) pooled cohorts from 1881 common era year (CE) versus age of death (years old; y). Curves show thin-plate spline fits (narrow confidence intervals in red dashed mostly overlay the black solid line fit). Vertical lines show ages with interpolated numbers of deaths 3/4 of numbers dying at the mode. In contrast to the pooled study with nine countries, the second-world-war direct-death mortality bump can hardly be seen at all with the male data. Source of raw data: Human Mortality Database (2019).

**Supplemental-FigS2 Modal death age summary statistics for seven countries (“EUM”)**. Female (+; red solid line: loess, grey area (hardly visible in b): standard error (SE)) and male (x; black dashed line: loess, grey area: SE) cohort modal death ages (years old; y) from (a) raw mortality (dx’) data; or (b) interpolated thin-plate splines; versus cohort birth date (common era year; CE) or date cohort defined as extinct (at 110 y). Interpolated pooled modal-death ages show a near-constant increase with female, but not male, cohorts. Source of raw data: Human Mortality Database (2019).

**Supplemental-FigS3 Cohort (a) median and (b) mean summary statistics for severn countries (“EUM”)**. Female (+; red dashed line: loess; grey area standard error (SE) hardly visible) and male (x; black solid line: loess; grey area SE hardly visible) cohort (a) median or (b) mean death ages (years old; y) from raw mortality (dx’) data, versus cohort birth date (common era year; CE) or date cohort defined as extinct (at 110 y). Raw male median-death ages show a slight decline over this period whereas females’ increase; raw male mean-death ages are almost constant. Source of raw data: Human Mortality Database (2019).

**Supplemental-FigS4 Cohort (a) median, and (b) mean, interpolated death ages for individual countries**. Female and male cohort loess-smoothed (a) median or (b) mean death ages (years old; y) from individual-country integrated thin-plate spline fits to mortality (dx’) data, versus cohort birth date (common era year; CE) or date cohort defined as extinct. Females (blue dot-dashed lines; italic labels); Males (red or black lines, upright labels): De, Denmark; Fi, Finland; Fr, France; Ic, Iceland; It, Italy; Ne, Netherlands; No, Norway; Sw, Sweden; Sz, Switzerland. Standard errors (grey). For both averages, female death-ages increased but five (red solid) countries male death-ages showed net decreases or were near-constant over the first 30 years. Source of raw data: Human Mortality Database (2019)

