## Supplementary figures and images for "Dramatic differences in male and female mortality trends for selected European cohorts over ∼20 years"

### Supplemental Fig S1

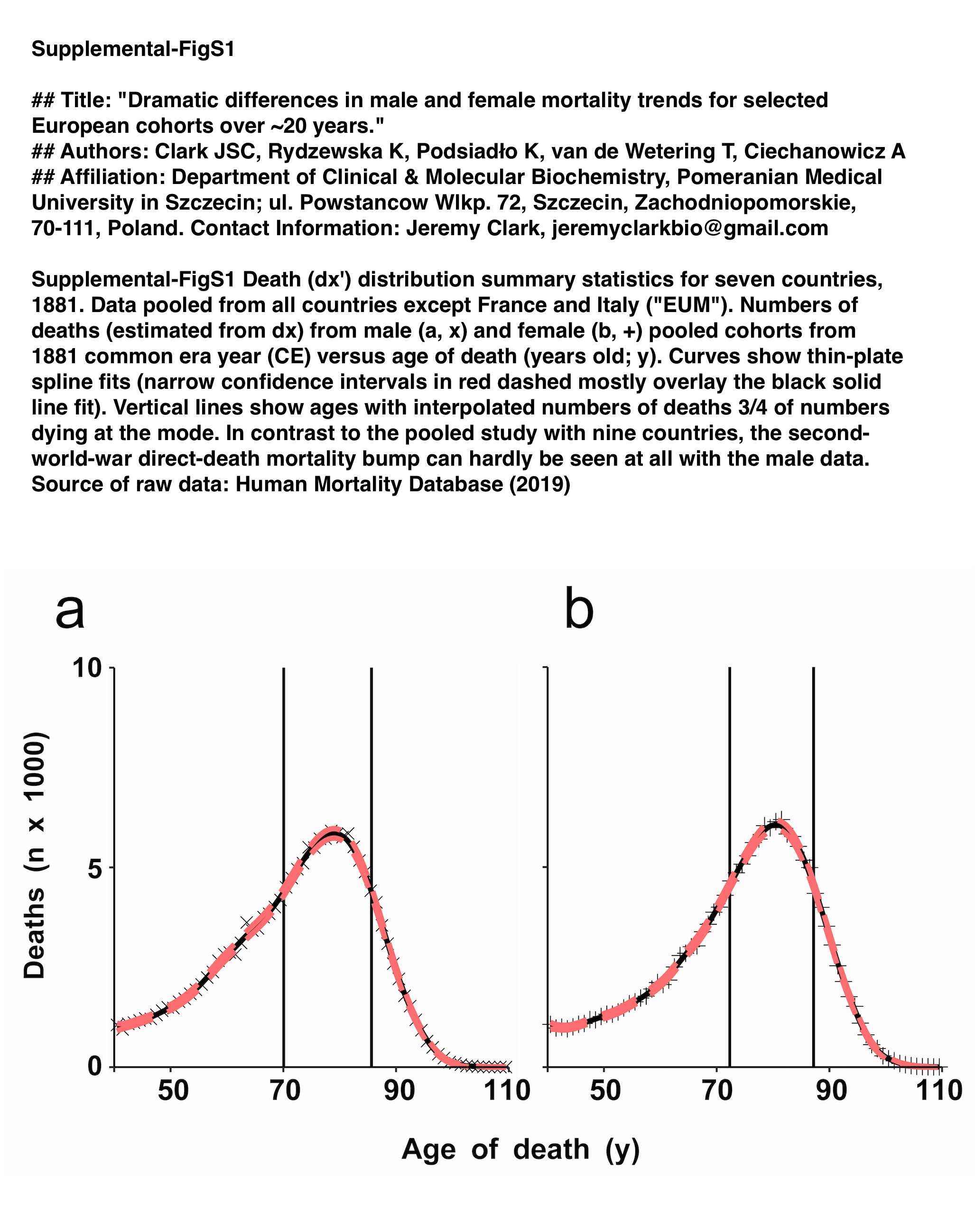

### Supplemental Fig S2

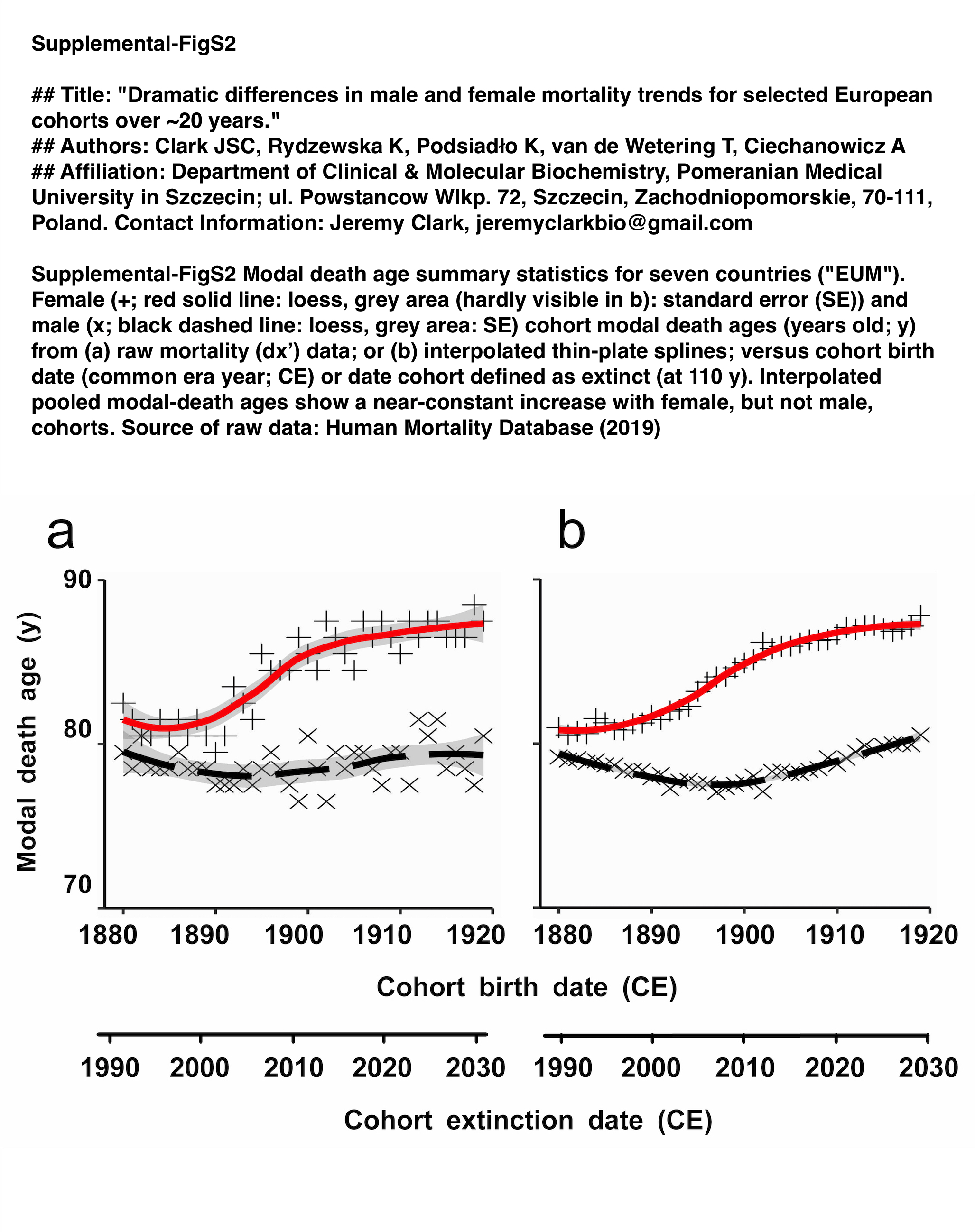

### Supplemental Fig S3

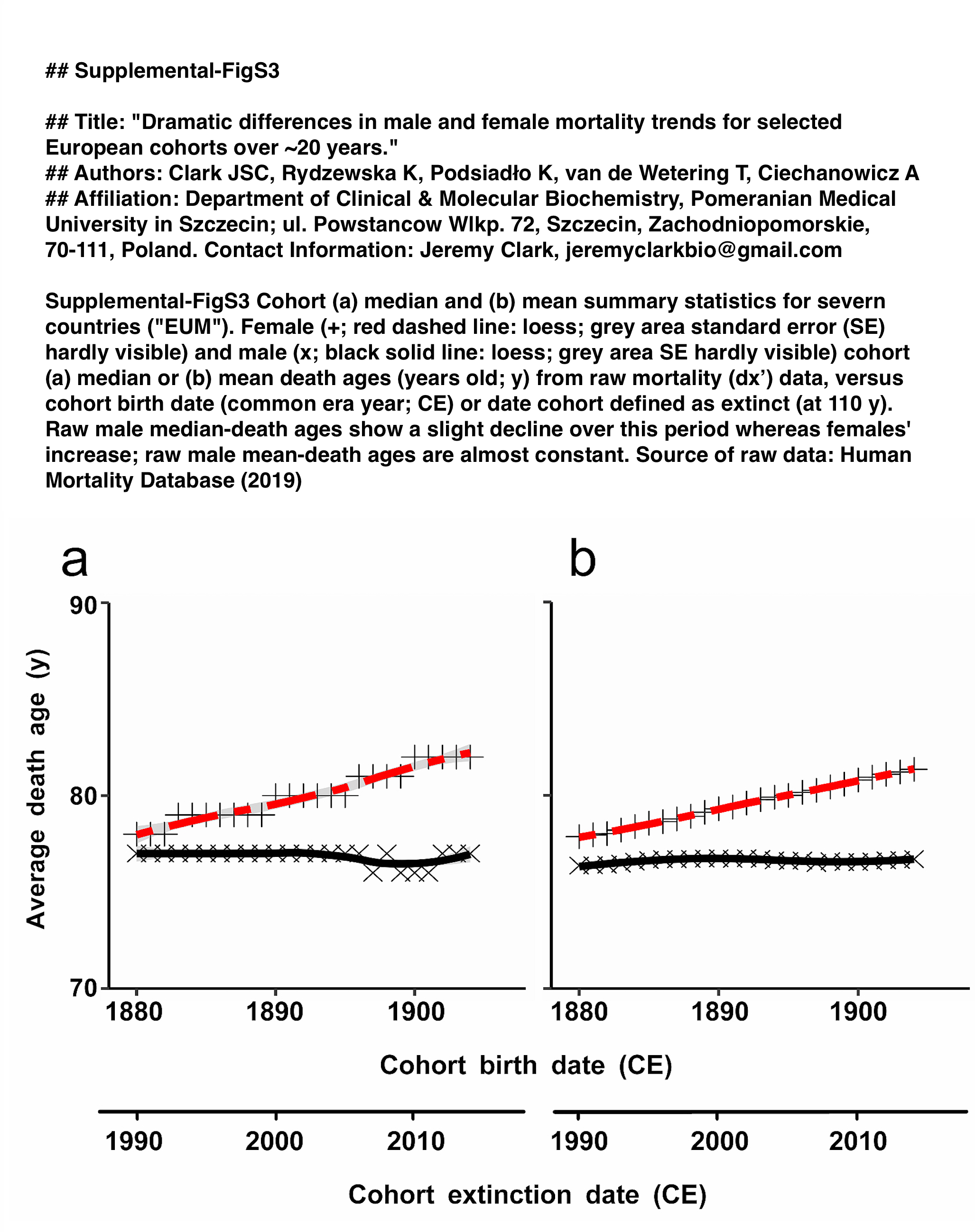

### Supplemental Fig S4

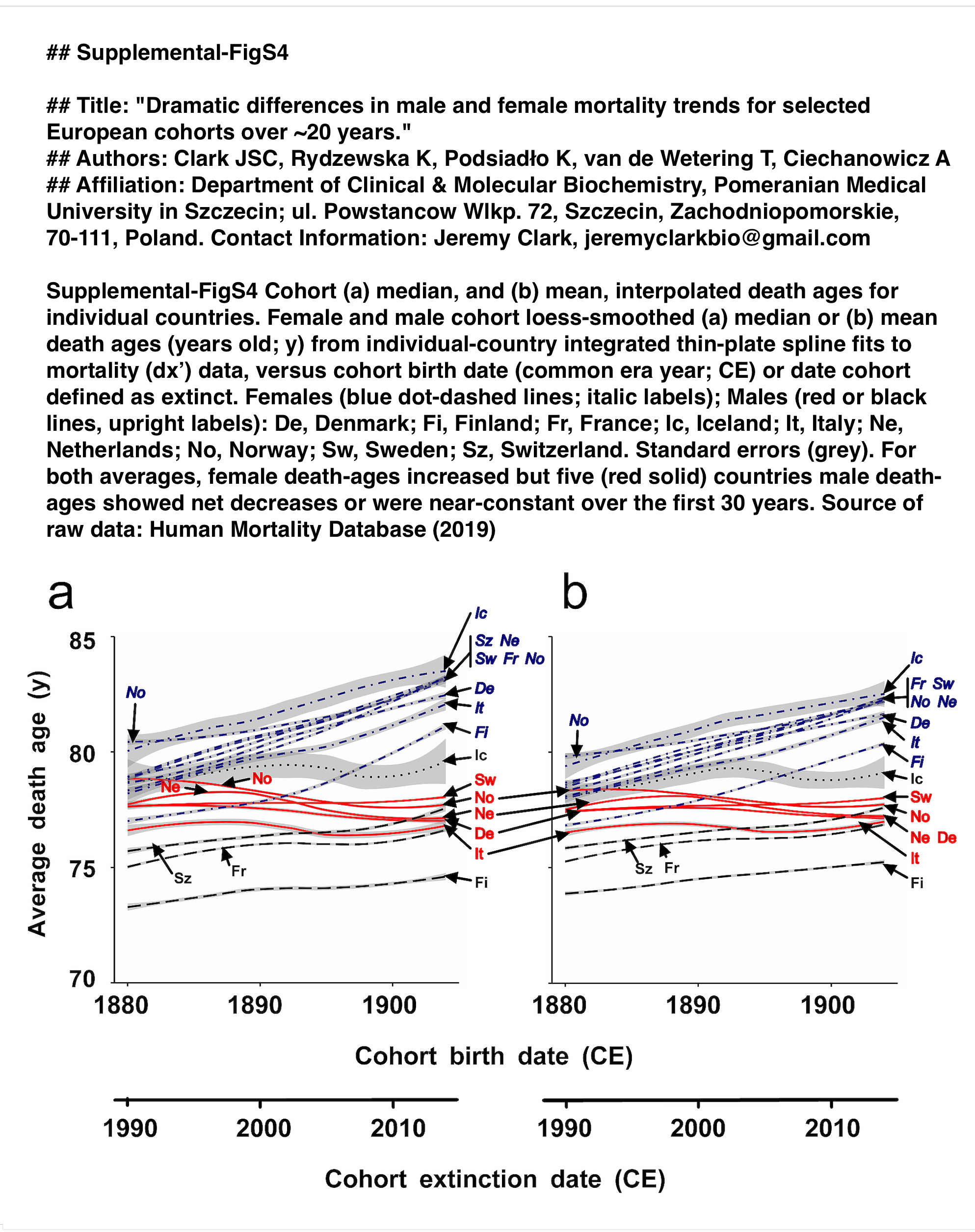
